# Effect of iris pigmentation of blue and brown eyed individuals with European ancestry on ability to see in low light conditions after a short-term dark adaption period

**DOI:** 10.1101/2024.01.17.576074

**Authors:** Faith Erin Cain, Kyoko Yamaguchi

**Affiliations:** School of Biological and Environmental Sciences, Faculty of Science, Liverpool John Moores University, Liverpool, Merseyside, United Kingdom

## Abstract

The effect of iris depigmentation on ability to see in low light conditions has not been thoroughly investigated as an adaptive advantage that could have contributed to the evolution and persistence of blue eyes in Europe. In this study 40 participants took part in a simple eye test in increasing luminance to examine if there was a difference in capacity to see in low light conditions between blue and brown-eyed individuals after a brief adaptation period. Blue eyed individuals were identified to have significantly better ability to see in lower lighting after a short adaption period than brown eyed individuals making it likely depigmented irises provide an adaptive advantage (p=0.046). Superior ability to see in low light conditions could be the result of increased straylight in depigmented irises, which in light luminance is disadvantageous but in low light conditions may provide an advantage. More research is needed to determine the specific association between melanin content and low-light visual acuity. Furthermore, more research is needed to establish that the improved capacity of blue-eyed individuals to see in low light settings seen in this study is attributable to iris pigmentation rather than corresponding pigmentation elsewhere.

## Introduction

The human iris or ‘rainbow membrane’ is considered the most complex tissue visible on the exterior of the human body with its variation in pattern and colour enabling real time identification [1, 2]. Eye colours include brown, intermediate and blue, however, eye colour variation is almost exclusively found in individuals of European descent [3]. The iris is made up of five cell layers. Melanin, an inert light-absorbing biopolymer pigment associated with human eye colour variation [4], is stored and synthesised by melanosomes within the melanocytes of the iris. Iris colour is determined by the quantity of melanin pigment granules in the anterior border layer and stroma [2, 5–8]. Pigmented irises appear brown due to their abundance of both forms of melanin; eumelanin (brown pigment) and pheomelanin (red-yellow pigment). Whereas blue (depigmented) irises contain very little melanin [4, 7]. Melanocyte number and melanosome size are not factors contributing to eye colour as they are constant across eye colours [7, 9].

The blue colour of depigmented irises is due to Tyndall Scattering in the relatively melanin free collagen fibrils of the stroma, that scatter the short blue wavelengths of light hitting the iris, demonstrating blue iris colour as a result of structural difference and not chemical composition, such as blue pigment [10]. There has been found to be no significant benefit to having a physician measure eye colour [11]. As such, iris colours can be classified according to systems such as Mackey, Wilkinson, Kearns and Hewitt (2011) nine-category grading system [12]. This system takes into consideration the pattern of pigmentation to assign irises into specific intermediate categories within three broader classification categories.

### Role of the iris

The iris plays a key role in human vision as it dictates the pupil size according to the light in an environment, governing the amount of light let into the eye [13]. Light let through the pupil is focused on the retina to provide vision. Barlow (1972) hypothesised that varying pupil size may facilitate rapid adaption to darkness by reducing rhodopsin bleaching at the retina [14, 15]. Alternatively, Campbell and Gregory (1960) suggest varying pupil size optimizes visual acuity by maximising light entering the pupil in low luminance and reducing loss of contrast by optical aberrations, straylight and diffraction in increased illumination [16]. As the iris is acutely adapted to control light entering the eye to provide visual acuity in a variation of illuminance levels, it would be logical that iris pigmentation has adapted to facilitate this function. However, iris pigmentation has been found to be independent to pupil size. Iris pigmentation was investigated as a factor affecting pupil size in a sample of 91 white subjects and it was concluded that iris pigmentation does not affect pupil size [13].

Iris depigmentation is only significantly observed in European populations with the highest frequency in the most northernly latitudes of Europe, which would have been part of the Eurasian tundra belt 10,000 years ago [3, 17]. Depigmentation of the iris to give rise to blue eye colouring has first appeared in the human population in Europe by point mutation in *HERC2* which reduced the activity of the *OCA2* promoter [18]. This A to G mutation at rs12913832 in *HERC2* is the main determinant of blue-eye colour [19] signature of selection has been found for the derived G allele that is associated with blue eye colour [20]. Despite the evidence for positive selection on blue iris colour in Europe, it is still unclear why blue iris colour emerged and persisted within the European population as there are contrasting arguments suggesting adaptive advantages of both pigmented and depigmented irises.

### Selection for iris depigmentation

In the Eurasian tundra belt, depigmented irises could have been selected for through rare-colour advantage among females from increased pressure of sexual selection due to a skewed operational sex ratio (OSR) caused by high mortality in males [17]. Additionally, individuals with depigmented irises could have been selected for through reduced susceptibility to Seasonal Affective Disorder (SAD), a recurring mood fluctuation with a seasonal pattern. The higher rates of SAD in brown-eyed individuals can result in suicidal levels of depression and reduced offspring production by social withdrawal, providing blue-eyed individuals with an advantage [21]. Selection for individuals with depigmented irises through rare colour advantage and reduced susceptibility to SAD could have contributed to the emergence and persistence of blue eye colour in Europe.

### Selection against iris depigmentation

Allowing more amount of light to enter the eye could act as a disadvantage by increasing susceptibility to disability glare. Blue/green-eyed individuals were found to have an increased straylight measure value compared to brown eyed individuals, increasing their disability glare [22]. Vos (2003) defined it as the “the masking effect caused by light scattered in the ocular media which produces a veiling luminance over the field of view” [23]. Disability glare could provide a selective disadvantage by impeding hunting abilities, particularly with light reflecting off snow in the Eurasian tundra. Furthermore, iris depigmentation could be disadvantageous through slower reaction times of motor response, to both auditory and visual stimuli, than individuals with pigmented irises. Landers *et al.* (1976) found mean reaction time to be 23 milliseconds faster in individuals with dark brown irises compared to those with blue irises in a sample of 48 Caucasian men and women [10]. Iris pigmentation is believed to indicate melanin in other parts of the body, for example Neuromelanin, which has a function in the speed of nerve impulses [10, 24]. This is thought to be the mechanism behind people with pigmented irises having faster reaction times than those with depigmented irises. Slower reaction times and increased disability glare are disadvantages of iris depigmentation which could contribute to selection against iris depigmentation.

The study by Bartholomew *et al.* (2016) found no significant difference in scotopic (night) visual acuity between individuals with blue-grey, green-hazel or brown-black after full dark adaption [25]. However, 52.7% of participants in the study were not of European ancestry but they did not statistically control for ancestry when testing the effect of iris colour on visually acuity or contrast sensitivity. It would be worth focusing on participants with European ancestry only to minimise the influence of genetic background. Also, their study examined visual acuity after complete adaption to low light conditions. Therefore, effect of iris pigmentation on visual acuity after a short adaption period warrants further investigation [26].

The aim of our study is to test a hypothesis that the ability to see in low light conditions was a selective pressure for depigmentation of irises in Europe by examining if pigmented and depigmented irises of living human differ in their capability to see in low light conditions after a short adaption period. This will be accomplished by measuring the capacity to read printed codes in steadily increasing light after a short adaption period. Improved visual acuity in low light settings after a short adaption period could provide a selective advantage when foraging between dark caves and daylight. The results of this study could help us to better understand the selective pressures acting upon the population of the Eurasian tundra belt during the emergence of depigmented irises.

## Methods

### Study Design

In this study, the ability of brown and blue eyes individuals to see in low light after a short dark adaptation period was compared by analysing participants’ ability to read codes in increasing light. The light level at which the participant could read the code was recorded and compared between blue and brown eyed individuals. Data was collected at the John Moore’s University Student Life Building between January and August 2022.

### Participants

Participants were recruited in line with the LJMU REC guidance following ethical approval by BESREC (approval reference number: 2021/BES/023) between January 2022 and July 2022. The study was concluded to be minimal risk as there was no threat to the psychological or physical wellbeing, values or dignity of the participant. Periods of time spent in darkness were kept minimal and intermittent to reduce potential distress to the participants. To maintain the investigation as minimal risk participants were not asked medical history, age, sex or other personal questions. Each participant was assigned a participant ID composed as a P and a number (P01 for example) to anonymise the data. Personally identifiable information was obtained in order to contact potential participants but no identifiable information was collected beyond this point and it is not possible to link any study data to participants. In advertisements and information sheets potential participants were asked only to apply if they meet the following criteria:

- Between 18-30 years old
- Of European descent
- Have blue or brown eye colour
- No history of laser eye surgery

Informed consent was attained from each participant by written consent form on the day of the test. A total of 40 individuals participated in this study. A larger sample size would have strengthened the results however there was not sufficient participant interest for this.

### Glasses/contact lens wearers

The effect of wearing glasses/contact lenses on ability to see in low light conditions was explored as an independent variable because there was an unequal distribution of glasses/contact lens wearers between the eye colours. It was noted if the participant wore glasses or made the investigator aware they wore contact lenses. During the test, these individuals were required to wear their glasses or contact lenses. The procedure for glasses/contact lens wearers was the same as the test for participants who did not wear contact lenses or glasses.

### Light Levels

The light levels were adjusted using the light box. The light box is a 29.5×25.0×18.0 cm cardboard box with 25 holes in the lid and contained 120 Lezonic string LED lights (Aaronic Tech Co., Ltd; Xiamen City, China). Each bulb was 3.6 watts. The light level was increased between each level by pulling another bulb up through the top of the box. The first light level was complete darkness with no lights exposed from the box in the light proof test room (40.55cm^3^).Light was prevented from leaving through the unoccupied holes in the box lid.

Lux was measured using the HoldPeak HP-881D Digital LUX Meter (HoldPeak instruments). The lux at the code at each light level was measured with the sensor of the luxmeter 132.5cm from the ground under the same conditions as the participant set up shown in Fig 1.

**Fig 1.**
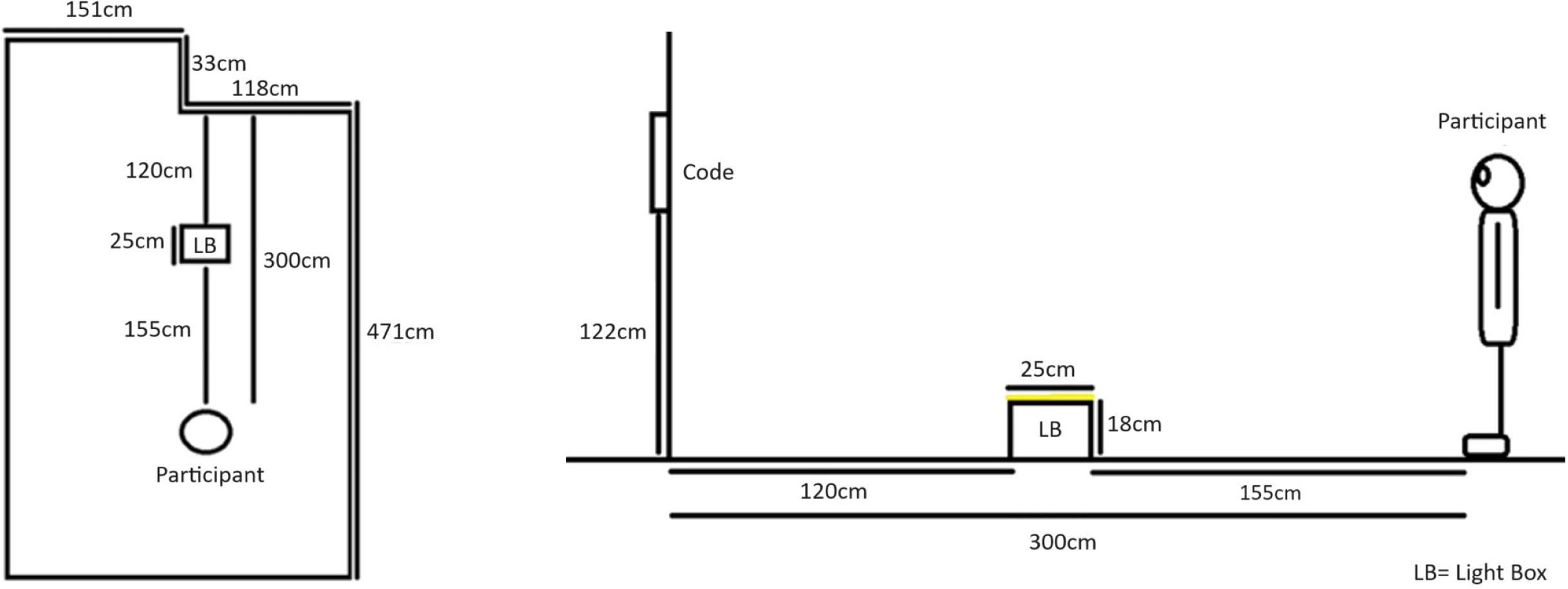
Participant set up for code reading test. A) birds eye view, B) side view: LB= light box.

Table 1 displays the average lux after 3 repetitions at the code based on the number of lights exposed from the box. A regular increase in lux was observed as more bulbs are added, each contributing an average of 0.068 lx (R^2^=0.9983, Fig 2).

**Fig 2.**
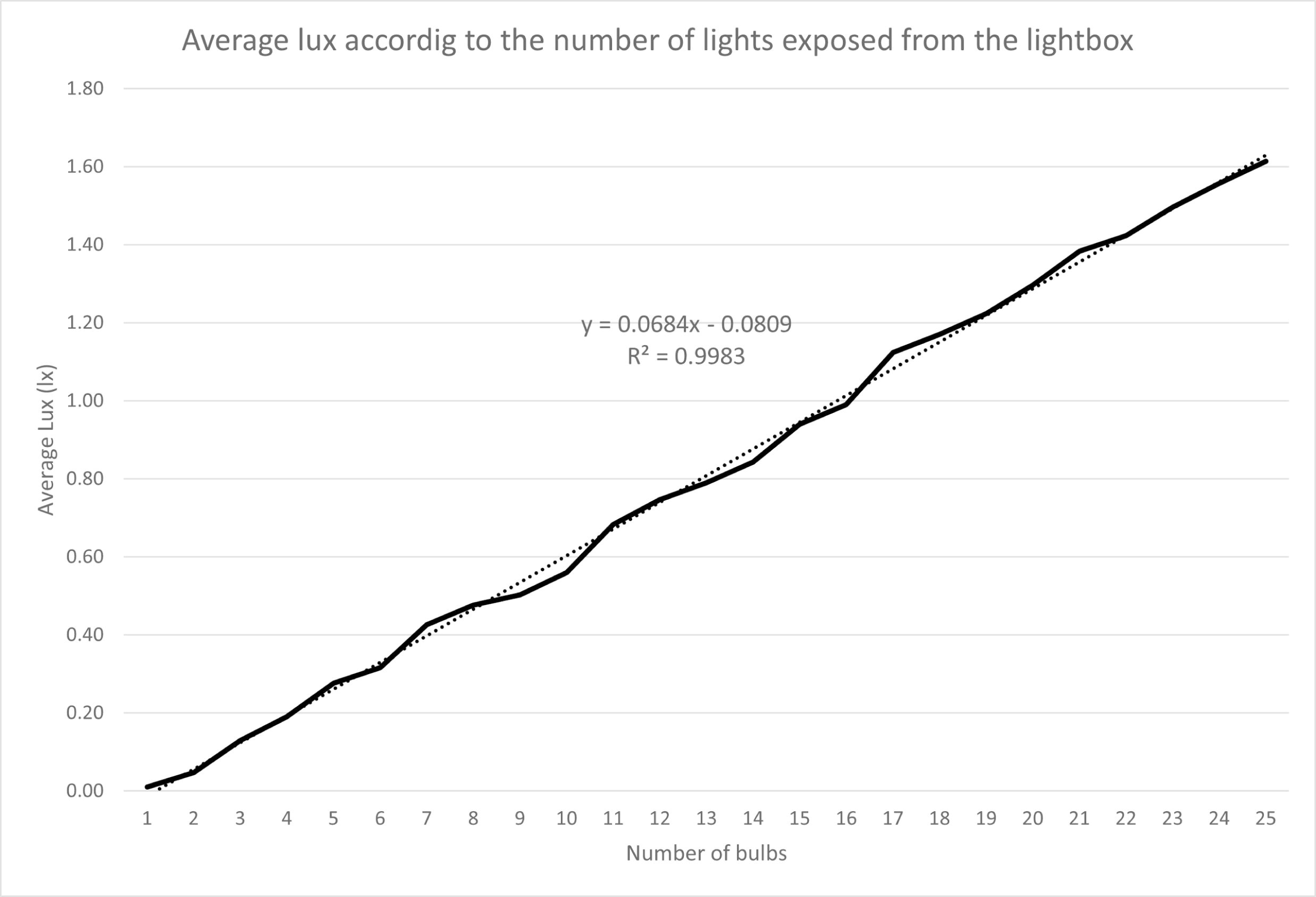
Average lux according to the number of lights exposed from the light box after 3 repetitions. The graph shows an increase in the average lux at the code with the number of lights exposed from the light box (y=0.0684x – 0.0809, R^2^=0.9983).

**Table 1.** Average Lux output of the light box measured on a vertical plane at the code after 3 repeats with an increasing number of bulbs.

| No. of Bulbs | Average Lux (lx) | Standard deviation | Lux per bulb (lx) |
| --- | --- | --- | --- |
| 0 | 0.01 | 0.00 | 0 |
| 1 | 0.05 | 0.01 | 0.04 |
| 2 | 0.13 | 0.00 | 0.08 |
| 3 | 0.19 | 0.00 | 0.06 |
| 4 | 0.28 | 0.01 | 0.09 |
| 5 | 0.32 | 0.00 | 0.04 |
| 6 | 0.43 | 0.01 | 0.11 |
| 7 | 0.48 | 0.01 | 0.05 |
| 8 | 0.50 | 0.00 | 0.03 |
| 9 | 0.56 | 0.01 | 0.06 |
| 10 | 0.68 | 0.01 | 0.12 |
| 11 | 0.75 | 0.01 | 0.06 |
| 12 | 0.79 | 0.01 | 0.04 |
| 13 | 0.84 | 0.01 | 0.05 |
| 14 | 0.94 | 0.00 | 0.10 |
| 15 | 0.99 | 0.02 | 0.05 |
| 16 | 1.12 | 0.01 | 0.13 |
| 17 | 1.17 | 0.00 | 0.05 |
| 18 | 1.22 | 0.01 | 0.05 |
| 19 | 1.30 | 0.01 | 0.07 |
| 20 | 1.38 | 0.01 | 0.09 |
| 21 | 1.42 | 0.03 | 0.04 |
| 22 | 1.50 | 0.01 | 0.07 |
| 23 | 1.56 | 0.01 | 0.06 |
| 24 | 1.61 | 0.01 | 0.06 |
| 25 | 1.67 | 0.01 | 0.06 |
| Weighted average lux per bulb: |  |  | 0.07 |
| SD lux per bulb: |  |  | 0.03 |

### Procedure

The process of the test is illustrated in Fig 3. Each test took approximately 30 minutes. During this time the participants had the test explained to them, signed consent forms, had their iris photographed and undertook the test.

**Fig 3.**
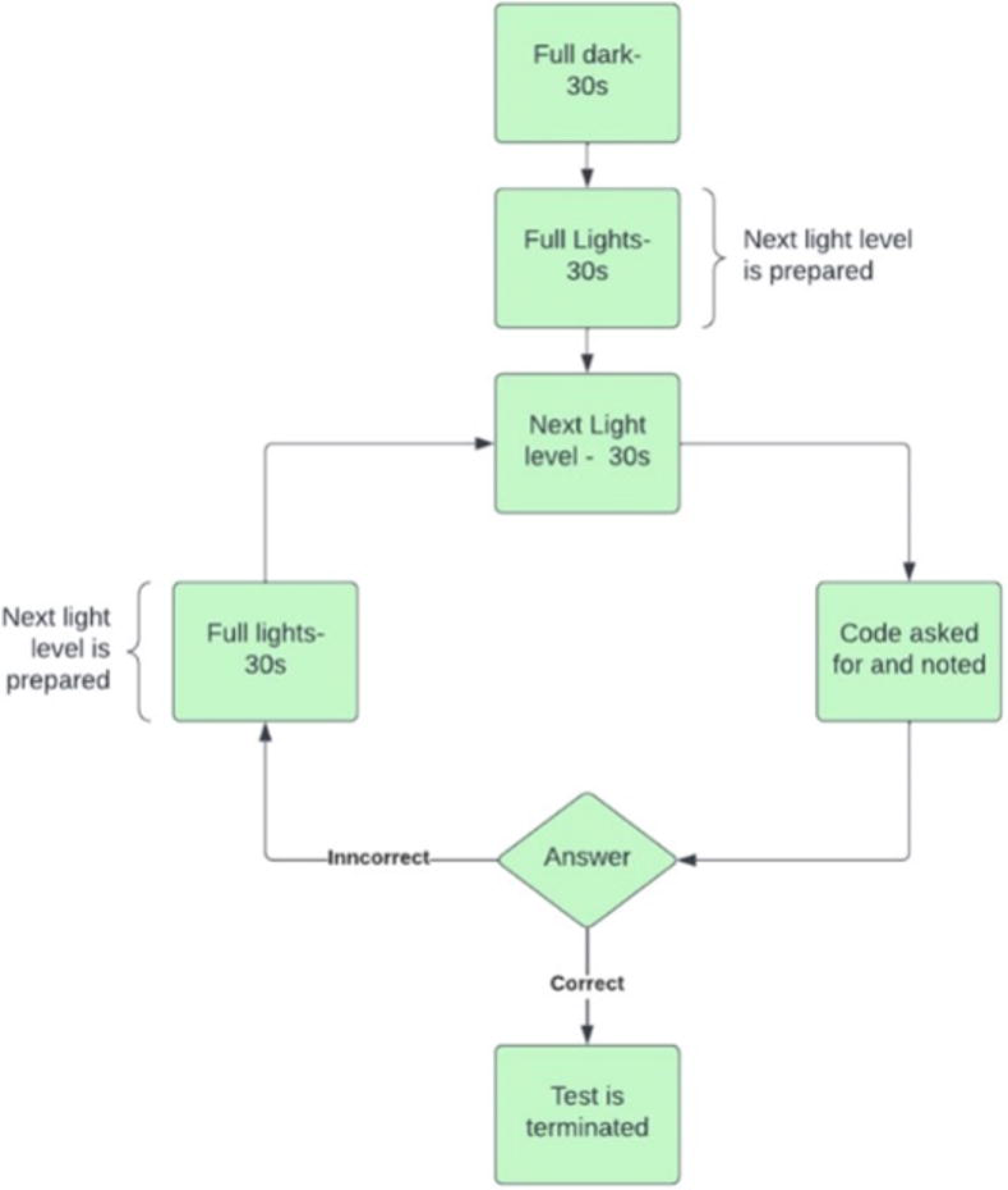
Diagram of the process during the participant test (s=seconds).

During the test, participants were asked to read a code on the adjacent wall 3 meters from them with the light box on the floor 120cm from the wall with the code, as illustrated by Fig 1. The distance from the code was chosen to be 3 meters following the Pelli-Robson contrast sensitivity chart test that uses the identification of letters to test visual acuity [27].

The test was comprised of 30 seconds of full light (with the main ceiling lights of the room on) followed by 30 seconds of the next light level (with the main ceiling lights of the room off) controlled by the light box. As complete rod and cone adaption to darkness after bright lights can take between 20 to 40 minutes, this study does not compare complete dark adaption between blue and brown eye colours [28–31].

Before the first light level each participant went through a preliminary 30 seconds of complete darkness followed by 30 seconds of full light. During the 30 seconds of full light the investigator would prepare the light box for the next light level by exposing another bulb from the top of the light box. Across the lux measurements the lights were exposed from the centre outward in the regular pattern shown in Fig 4. During the following 30 seconds of the next light level the investigator secured the code 122cm off the ground and the participant was asked to look ahead at the code and particularly avoid looking at the light box to avoid disability glare. When prompted by the investigator at the end of the 30 seconds the participant would be asked to read what they could see of the code which was noted by the investigator. The code was then removed from the wall before starting the next 30 seconds of full light to prepare for the next light level. Also, during the 30 seconds of full light, note was taken if the participant had been able to correctly read the code in the pervious light level. Once the participant had correctly read the code the test was continued for three more light levels before the participant was informed the test was complete. The variable used in this study is the light level score which corresponds to the number of exposed lights at the light level they were first able to read the code.

**Fig 4.**
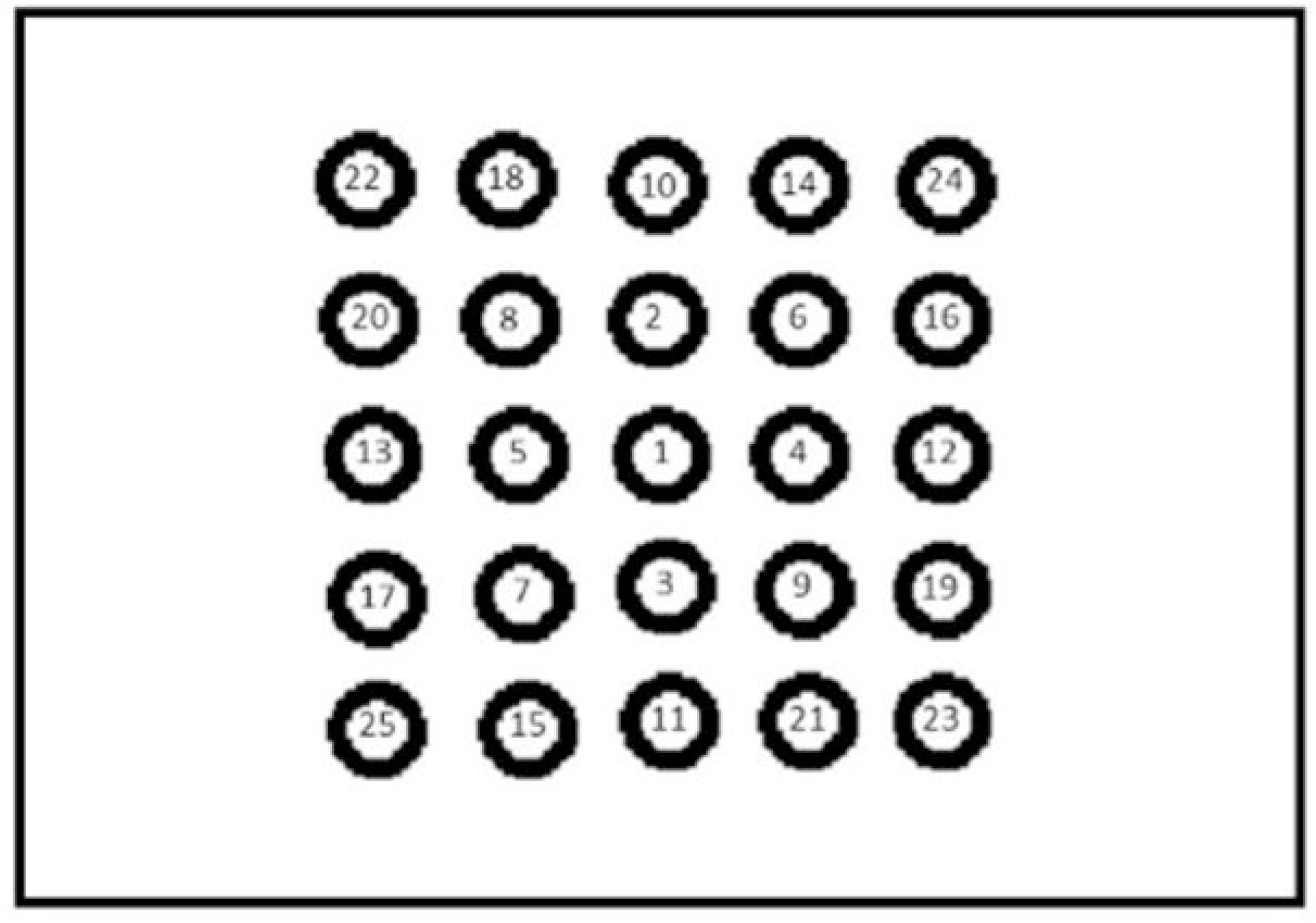
Diagram showing the pattern of bulb exposure from the top of the light box.

### Codes

Each light level had a specific code for the participants to read. Each code was composed of five randomly generated capital letters in Calibri font size 190 in black (RGB: 0,0,0). The letters were horizontal on A4 dark grey paper (RGB: 51,51,51). In order to get the code correct the participant had to read all five letters in order.

### Iris Classification

Irises were photographed with the participant stood against a white wall. Photographs were taken on an iPhone XR (Apple Inc.) back camera using flash from approximately 6 cm away from the eye. To remove subjectivity from determining iris colour, six colour samples were taken from the iris photos and their RGB values were analysed using the eyedropper tool of Photoshop, version 3.0 (Adobe Systems, 2021). Average RGB values for the peripupillary ring area and periphery iris area (overall eye colour) were generated from three colour samples from each area in a triangle shape (Fig 5). The RGB value (Red, Green or Blue) present in the highest quantity reflected the unobjective colour of that part of the iris. Brown eye colour was represented by red, blue eye colour by blue, and green intermediate colours by green. Irises were categorised by the colours of these areas according to Mackey *et al.*’s (2011) Iris colour classification grading system [12].

**Fig 5.**
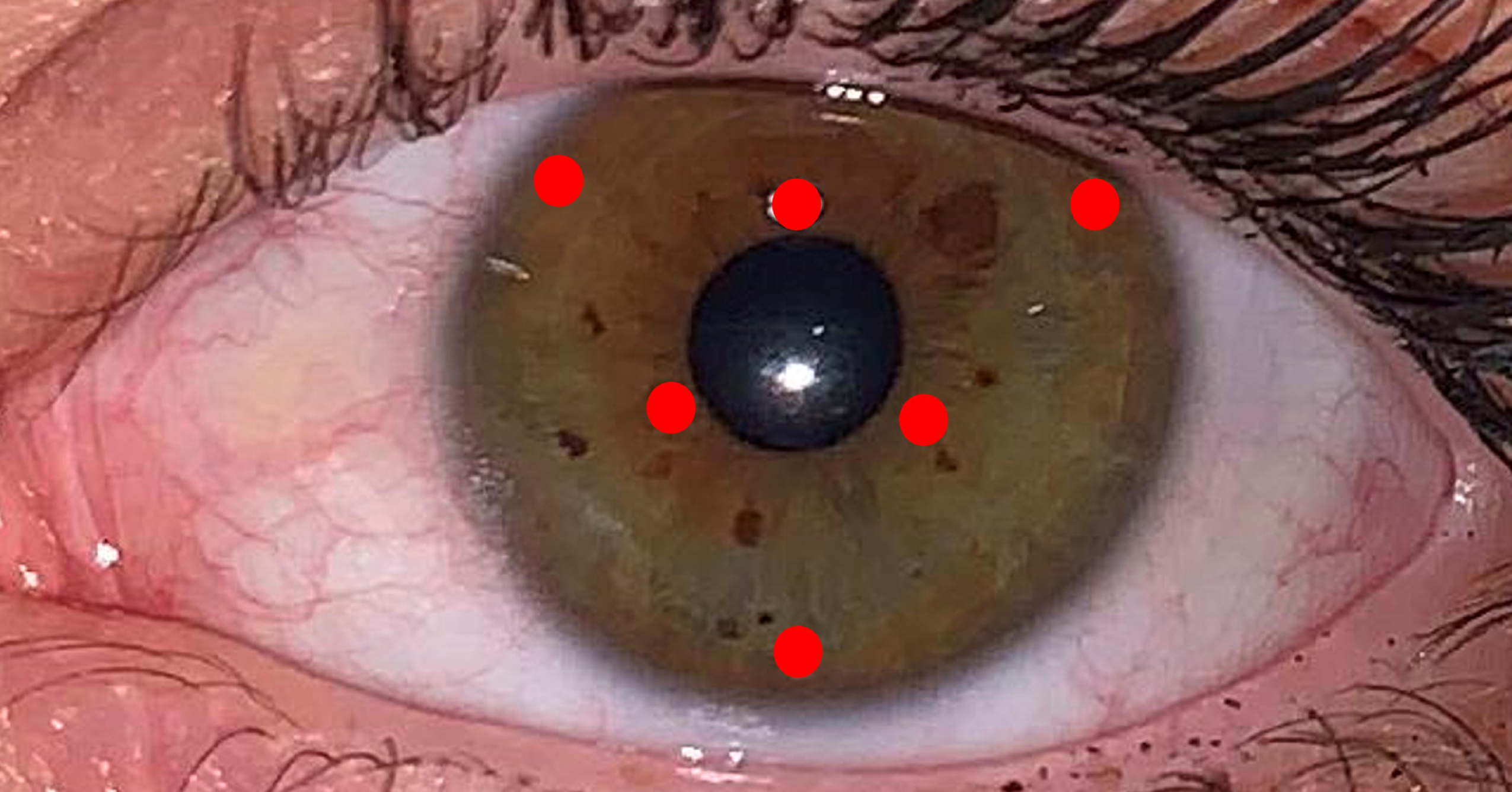
The six colour sampling points on the iris. The RGB values of the six colour samples from the iris photos were analysed with the eyedropper tool of Photoshop, version 3.0 (Adobe Systems).

### Statistical analysis

Descriptive statistics of the participants score were calculated using Microsoft Excel version 2108.

Outliers were identified within each eye colour group using the outlier labelling rule with the standard 1.5 multiplier suggest by Tukey (1977) [32]. The outlier labelling rule was also used with a 2.2 multiplier as recommended by Hoaglin and Iglewicz (1987) for more accurate outlier labelling in smaller sample sizes [33].

Due to small sample size nonparametric tests were employed. SPSS Statistics version 27 (SPSS, IBM) was first used to conduct a test of homogeneity to ensure the data fits the assumptions of Mann–Whitney U; same distribution of data across blue and brown eye colour. A Mann–Whitney U test was conducted to compare differences in light level scores between blue and brown eye colours, again using SPSS. A second Mann–Whitney U test was conducted to compare differences in light level scores between glasses and contact lens wearers and non-wearers.

## Results

### Iris colour analysis

The iris of one participant did not fit into blue or brown eye colour groups by Mackey *et al.’s* (2011) Iris colour classification grading system [11]. This participant was excluded from further analysis. Therefore, self-reported eye colour was 97.5% (39/40) consistent with colour analysis. The results of colour analysis are displayed in Table 2. The original data are available in S1. Within the sample used in this study, 36% (14/39) of participants had brown eyes and 64% (25/39) had blue eyes. The colour category with lowest and highest mean light level score was light blue and brown with peripheral green respectively.

**Table 2.** Eye colour classification category according to Mackey et al.’s (2011) and mean light level score (number of bulbs) of participants.

|  | Eye colour classification category | Quantity | Mean<br>Light<br>level<br>score |
| --- | --- | --- | --- |
| Blue (25) | Light Blue | 12 | 10.08 |
|  | Dark Blue | 8 | 10.5 |
|  | Blue with peripuillary brown | 5 | 10.2 |
| Brown (14) | Brown with peripheral green | 2 | 13 |
|  | Light Brown | 5 | 12 |
|  | Dark Brown | 7 | 11.71 |

### Light level score according to eye colour

Descriptive statistics of the light level score according to eye colour are found in Table 3. The mean light level score of brown eyed individuals was 1.76 light levels (0.12lx) greater than blue eyed individuals and 1.13 (0.08lx) greater than the combined mean. Blue eyed light level scores ranged from 7 to 17, which is corresponding to an average lux of 0.43lx to 1.12 lx at the code. Brown eyed light level scores ranged from 8 to 19, which is corresponding to an average lux of 0.48 lx to 1.22 lx at the code.

**Table 3.** Descriptive statistic of light level scores (number of bulbs) of blue, brown and combined groupings.

|  | Sample |  |  |
| --- | --- | --- | --- |
|  | Blue | Brown | Combined |
| Mean | 10.24 | 12 | 10.87 |
| Standard Error | 0.55 | 0.8 | 0.47 |
| Median | 10 | 11.5 | 10 |
| Mode | 9 | 11 | 9 |
| Standard Deviation | 2.76 | 2.99 | 2.93 |
| Range | 10 | 11 | 12 |
| Minimum | 7 | 8 | 7 |
| Maximum | 17 | 19 | 19 |
| Count | 25 | 14 | 39 |
| Percentage | 64 | 36 | 100 |

The light level scores of P26 (Score=17) and P27 (Score=17) were identified as outliers using the outlier labelling rule with a multiplier of 1.5. However, they were not identified as outliers using a multiplier of 2.2 that is suitable for a small sample size. Using a 1.5 or 2.2 multiplier, no brown-eyed outliers were found.

The test of homogeneity of variance showed equal distribution demonstrating the data fit the assumptions of the Mann-Whitney U test (p=0.651). The Mann-Whitney U test revealed a significant difference in light level score between brown eyed individuals (Mean=12 bulbs/0.82lx) and blue-eyed individuals (Mean= 10.24 bulbs/ 0.70lx; U=107.5, *n*_1_=14, *n*_2_=25, p=0.046). The results of the Mann-Whitney U test remained significant after the exclusion of the two outliers (U=81.5, p=0.012). Distribution of participants light level score according to eye colour can be seen in Fig 6.

**Fig 6.**
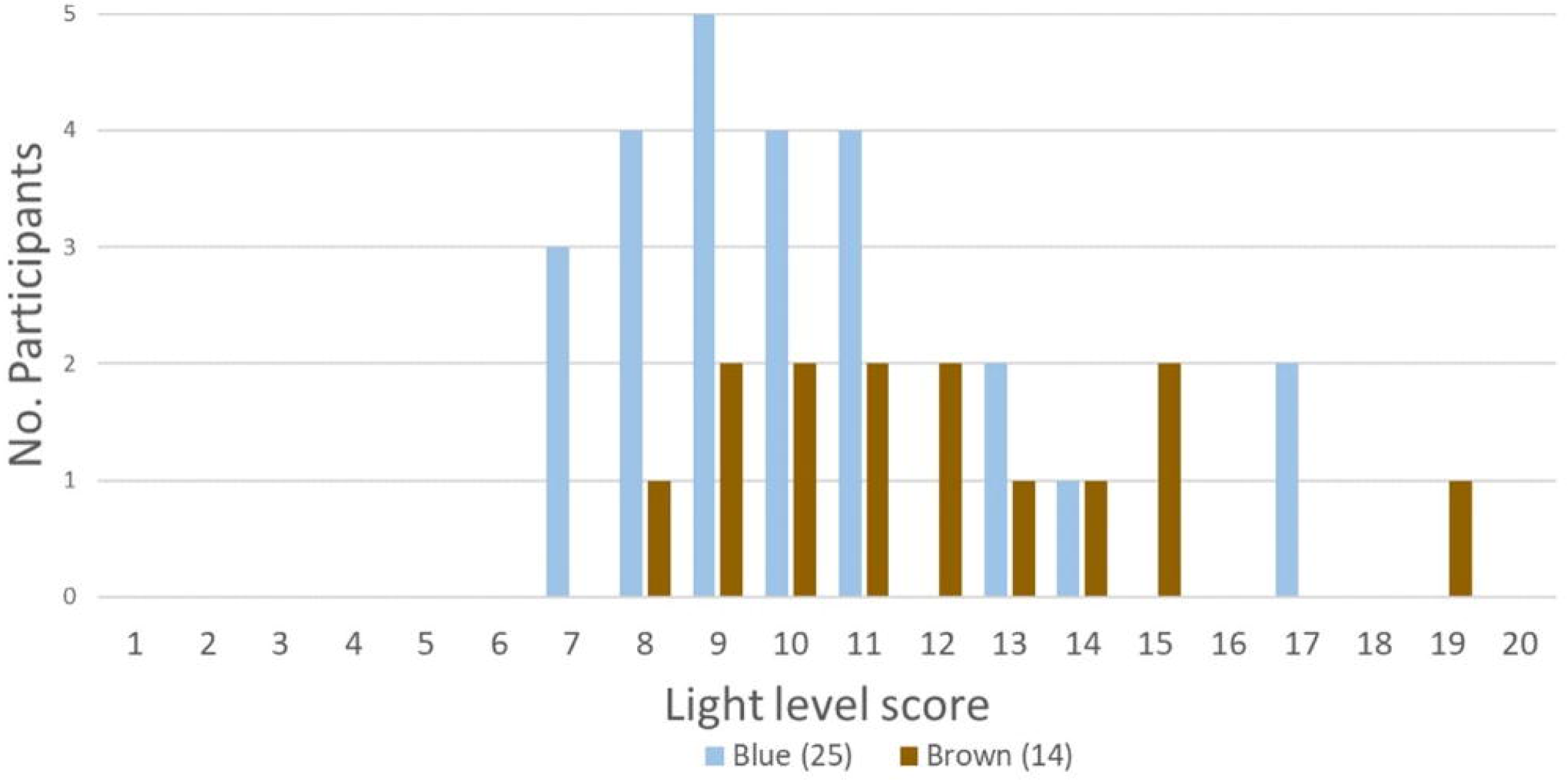
Distribution of participants light level score according to eye colour.

Light level score according to whether glasses/contact lenses are worn Within this study the percentage of glasses wearers was 1.43% higher in brown eyed individuals (3/14 = 21.43%) than brown eyed individuals (5/25 = 20.0%) (S1 Table).

Descriptive statistics of the light level scores according to whether glasses/contact lenses are worn are found in Table 4. The mean light level score of wearers was 0.8 (0.05lx) light levels greater than non-wearers and 0.63 (0.04lx) greater than the combined mean. Non-wearers light level scores ranged from 7 to 19, which is corresponding to an average lux of 0.43lx to 1.22lx at the code. Wearers light level scores ranged from 8 to 17, which is corresponding to an average lux of 0.48lx to 1.12lx at the code.

**Table 4.**
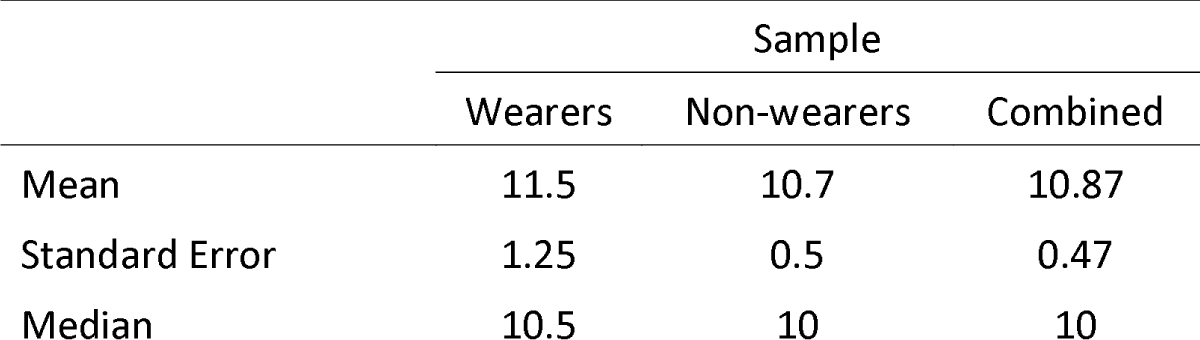
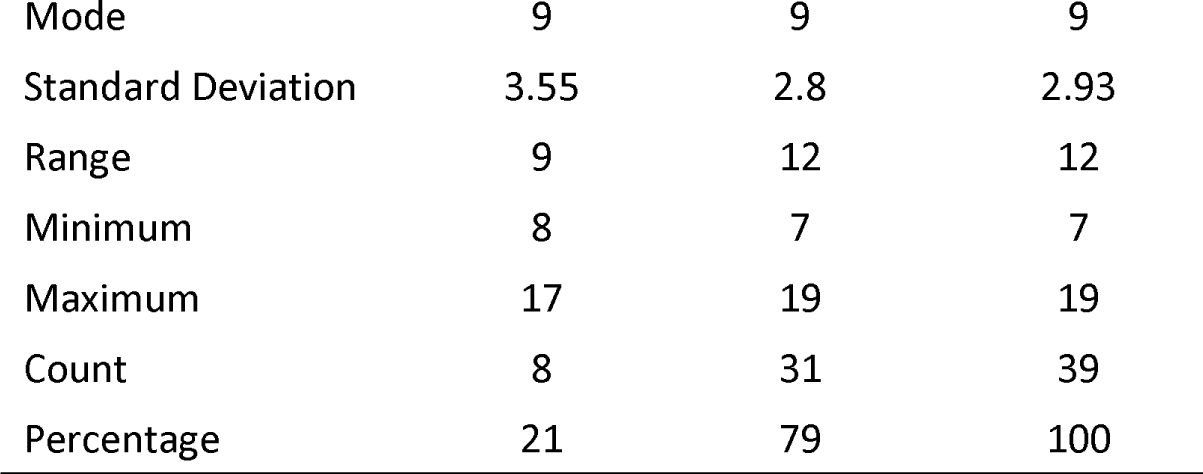
Descriptive statistic of light level scores of glasses/contact lens wearers, non-glasses/contact lens wearers and combined.

A Man-Whitney U test revealed insignificant differences in the light level score of glasses/contact wearers (Mean=11.50 bulbs/ 0.79lx) and non-wearers (Mean=10.70 bulbs/ 0.73lx; U=111.5 p=0.661). This justified the inclusion of glasses and contact lens wearers in the investigation into iris pigmentation. The test of homogeneity of variance showed equal distribution demonstrating the data fit the assumptions of the Mann-Whitney U test (p=0.651).

## Discussion

The results of this study reveal that blue-eyed individuals can read codes in significantly less light than brown-eyed individuals, with blue-eyed individuals reading the code with an average of 10.24 (0.70lx) light exposed from the light box compared to 12 (0.82lx) lights exposed for brown-eyed individuals (p=0.046). This infers blue-eyed individuals have an ability to see in low light conditions that is superior to that of brown eyed individuals.

Providing blue eyed individuals with a selective advantage could be the basis of depigmented irises become a prominent trait within the European population.

### Increased straylight in blue irises as a potential advantage in low light conditions

As previously stated, straylight is thought to cause a visual disadvantage since it reduces contrast and commonly manifests itself as disability glare [22, 34, 35]. However, this may not be the case in low light, as demonstrated by our finding that blue-eyed people have a better ability to see in low light after a short adaption period. Straylight is dependent on iris pigmentation as Ijspeert *et al.* (1990) found blue eyes to have a mean straylight measure of 0.949 across three glare angles compared to brown eyes with a straylight measure of 0.858 [22]. In fact, brown irises transmit approximately 100 times less light than blue irises [35]. It could be hypothesised that, in brown eyed individuals, melanin in the anterior border layer and stroma of the iris absorbs straylight that would otherwise pass through the pupil and cast a veil of light on the retina. For blue eyed individuals, in low light conditions after a short adaption period, this veil of light contributes enough luminance to provide blue-eyed individuals with a visual advantage to make out shapes. This phenomenon is demonstrated by blue-eyed individuals being able to read codes in less light than brown eyed individuals in our study. We demonstrated that illumination is the limiting factor of visual acuity in low light conditions, where blue eyed individuals have the advantage. However, contrast sensitivity, hindered by straylight, is the limiting factor which provides brown eyed individuals with the visual advantage in average to high light conditions [36].

Sturm and Larsson (2009) speculated that iris pigmentation influences visual acuity in low light conditions [8]. The findings of our study are contradictory to this as blue and brown-eyed individuals had significantly different light level scores (p=0.046). It has been reported that brown irises transmit significantly less straylight than blue irises and are less susceptible to disability glare [22, 26, 35]. Since our study hypothesises that increased light level score (needing more light to read the code) is the product of decreased straylight, the findings of our study are consistent with the findings of the previous studies [22, 26, 35].

### Influence of glasses and contact lens

To justify the inclusion of glasses/contact lens wearers in the sample comparing light level score between brown and blue-eyed individuals, the light level score between glasses/contact lens wearers and non-wearers was investigated. Because the number of individuals who use glasses or contact lenses was not evenly distributed between blue and brown eyed individuals, this investigation was necessary (Blue=25% and Brown=27.27%). If wearing glasses/contact lenses affects light level score, an unequal proportion of glasses/contact lens wearers may cause one eye colour to be influenced more than the other. According to Van Der Meulen *et al.* (2010), rigid contact lenses cause more straylight during and after use. According to the concept that more straylight leads to a lower light level score, wearing glasses or contact lenses could lower the light level score [37]. Also, glasses and contact lenses have blue light filters which could affect light level score, which was not controlled in this study. However, a study by Hammond (2015) suggested no effect of blue light filtering lenses on visual acuity [38]. Also, our study shows no significant difference (p=0.661) in light level score between glasses and contact lens wearers and non-wearers. In light of this, the current study warrants the inclusion of glasses/contact lens wearers in the investigation into ability to see in low light conditions between blue and brown eye colour.

### Other factors

As this is a preliminary study with a small sample size, we have only recruited individuals with blue or brown eye colours. In the future, a study with a larger sample including intermediate eye colours and measuring melanin content will be able to provide further insight into the direct relationship between iris pigmentation and ability to see in low light conditions.

Apart from iris pigmentation, other factors could have contributed more to visual acuity. For example, diet has been linked to vision in darkness, particularly the effect of malnutrition and vitamin A deficiency [39]. It is not unexpected that vitamin A plays a role in night vision as it is a precursor of rhodopsin; a photopigment found in rods within the retina [40]. A questionnaire on diet could be added to the procedure to control for the effect of dietary factors. Further study is required to isolate pigmentation of the iris as the source of variation in ability to see in low light conditions observed in this study.

## Conclusion

The findings of this study show that after a short adaptation period, blue eyed individuals have greater capacity to see in low light conditions than brown eyed individuals. The advantage of greater visual acuity in low light conditions after a short adaption period could have been the basis of the emergence and persistence of blue eye colour within the European population. Through comparison with other studies comparing blue and brown irises, increased visibility in low light conditions could be the product of increased straylight in blue irises which casts a veil of light over the retina. Further study is need to fully understand the relationship between the iris pigmentation and ability to see in low light conditions.

## Supporting Information

**S1 Table. Recorded number of light bulbs required to read a code by participants with their eye colours and glasses wearing status.** The ‘specific eye colour’ was determined using a photograph of each participant’s iris by Mackey et al.’s (2011) Iris colour classification grading system [12]

