## Supplementary material for "Effect of iris pigmentation of blue and brown eyed individuals with European ancestry on ability to see in low light conditions after a short-term dark adaption period": S1 Table

S1 Table. Recorded number of light bulbs required to read a code by participants with their eye colours and glasses wearing status.

| No. light bulbs required to read a code | Eye Colour | Specific eye colour | Glasses |
|---|---|---|---|
| 7 | Blue | Light Blue | No |
| 7 | Blue | Light Blue | No |
| 8 | Blue | Light Blue | No |
| 9 | Blue | Light Blue | No |
| 9 | Blue | Light Blue | No |
| 9 | Blue | Light Blue | No |
| 10 | Blue | Light Blue | Yes |
| 10 | Blue | Light Blue | No |
| 11 | Blue | Light Blue | Yes |
| 11 | Blue | Light Blue | No |
| 13 | Blue | Light Blue | No |
| 17 | Blue | Light Blue | Yes |
| 8 | Blue | Dark Blue | No |
| 8 | Blue | Dark Blue | No |
| 8 | Blue | Dark Blue | No |
| 9 | Blue | Dark Blue | Yes |
| 10 | Blue | Dark Blue | No |
| 11 | Blue | Dark Blue | No |
| 13 | Blue | Dark Blue | No |
| 17 | Blue | Dark blue | Yes |
| 7 | Blue | Blue with Peripullary Brown | No |
| 9 | Blue | Blue with Peripullary Brown | No |
| 10 | Blue | Blue with Peripullary Brown | No |
| 11 | Blue | Blue with Peripullary Brown | No |
| 14 | Blue | Blue with Peripullary Brown | No |
| 12 | Brown | Central brown and peripheral green | No |
| 14 | Brown | Central brown and peripheral green | No |
| 8 | Brown | Light Brown | Yes |
| 9 | Brown | Light Brown | No |
| 11 | Brown | Light Brown | Yes |
| 13 | Brown | Light Brown | No |
| 19 | Brown | Light Brown | No |
| 9 | Brown | Dark Brown | Yes |
| 10 | Brown | Dark Brown | No |
| 10 | Brown | Dark Brown | No |
| 11 | Brown | Dark Brown | No |
| 12 | Brown | Dark Brown | No |
| 15 | Brown | Dark Brown | No |
| 15 | Brown | Dark Brown | No |
